# Holotype genome of the lesula provides insights into demography and evolution of a threatened primate lineage

**DOI:** 10.1101/2025.06.05.657981

**Authors:** Axel Jensen, Emma R. Horton, Koko Bisimwa, Kate M. Detwiler, Katerina Guschanski

## Abstract

The rapid development of genome sequencing techniques in recent decades has revolutionized the field of evolutionary biology, facilitating the study of adaptation and speciation at the genome level. Genomic data has also become a cornerstone in conservation management, allowing inferences of population demography and genetic diversity. Here, we sequenced the genome of the holotype specimen of the elusive lesula (*Cercopithecus lomamiensis*), a recently described member of the guenons (tribe Cercopithecini) endemic to the Democratic Republic of the Congo. Using published and novel genomic data, we explore the evolutionary and demographic history of *C. lomamiensis* and its sister species *C. hamlyni*, the only two members of the *hamlyni* species group. We estimate that the two species split ca. 3-4 million years ago, and find that they both show high genetic diversity despite being listed as vulnerable on the IUCN Red List. We identify signatures of positive selection in the *hamlyni* group in genes involved in, e.g., pelage coloration and immune functions, as well as skeletal morphology and locomotor behavior, potentially related to their terrestrial lifestyle which stands out among the otherwise arboreal *Cercopithecus* genus. We specifically explored whether introgression from more distantly related terrestrial guenons was involved in the evolution of terrestriality in the *hamlyni* group. However, molecular convergence with other terrestrial species was low, suggesting that putative terrestrial adaptations occurred largely independently. By sequencing the *C. lomamiensis* genome, this study provides insights into the demography and evolutionary history in a threatened primate lineage, with relevance to conservation management.

## Introduction

With the emergence of available and affordable sequencing technologies, whole genome sequencing data has become an essential tool in conservation management (Formenti et al., 2022). Neutral genomic markers can provide insights into demographic processes such as inbreeding and effective population size, and variation in coding regions can be utilized to explore lineage-specific adaptations.

Whole-genome sequencing data complements field-based research of species biology, allowing for inferences that are difficult or impossible to obtain through direct observations. In poorly known taxa that are difficult to study in the field, genomic analyses may thus be vital to aid conservation management, as well as taxonomic classification. Consequently, sequencing the genome of type specimens is increasingly acknowledged as an important complement to the species description (Federhen, 2015; Renner et al., 2024). In addition to its importance for proper taxonomic and conservation assessment, sequencing from type material serves as a valuable reference for future genomics studies.

*Cercopithecus lomamiensis* (common name: lesula) is an elusive primate species endemic to the Democratic Republic of the Congo (DRC), which was unknown to the scientific community until 2012 (Hart et al., 2012). It has been a protected species in the DRC since 2016, following the establishment of the Lomami National Park in the core of its distribution range. The scientific discovery of *C. lomamiensis* was among the key elements that led to the park’s establishment, with the species’ distinctive face becoming the official logo for the park. Based on appearance, vocalizations, ecology and molecular marker data, Hart et al. (2012) placed *C. lomamiensis* as a distinct sister species to *C. hamlyni* (owl-faced monkey), from which it is geographically separated by the Congo and Lomami rivers and their interfluvial region (Figure 1A). Together, *C. lomamiensis* and *C. hamlyni* form the *hamlyni* species group (Figure 1B), as members of the guenons (tribe *Cercopithecini*), a diverse group of African primates (Grubb et al., 2003). Although *C. hamlyni* has a relatively large distribution range in eastern DRC (Figure 1A; Hart & Maisels, 2020), the *hamlyni* species group remains understudied. Both species, *C. lomamiensis* and *C. hamlyni*, albeit locally abundant, are listed as vulnerable on the IUCN Red List, primarily due to habitat loss and hunting (Detwiler & Hart, 2020; Fournier, Graefen, et al., 2023; Hart & Maisels, 2020).

**Figure 1.**
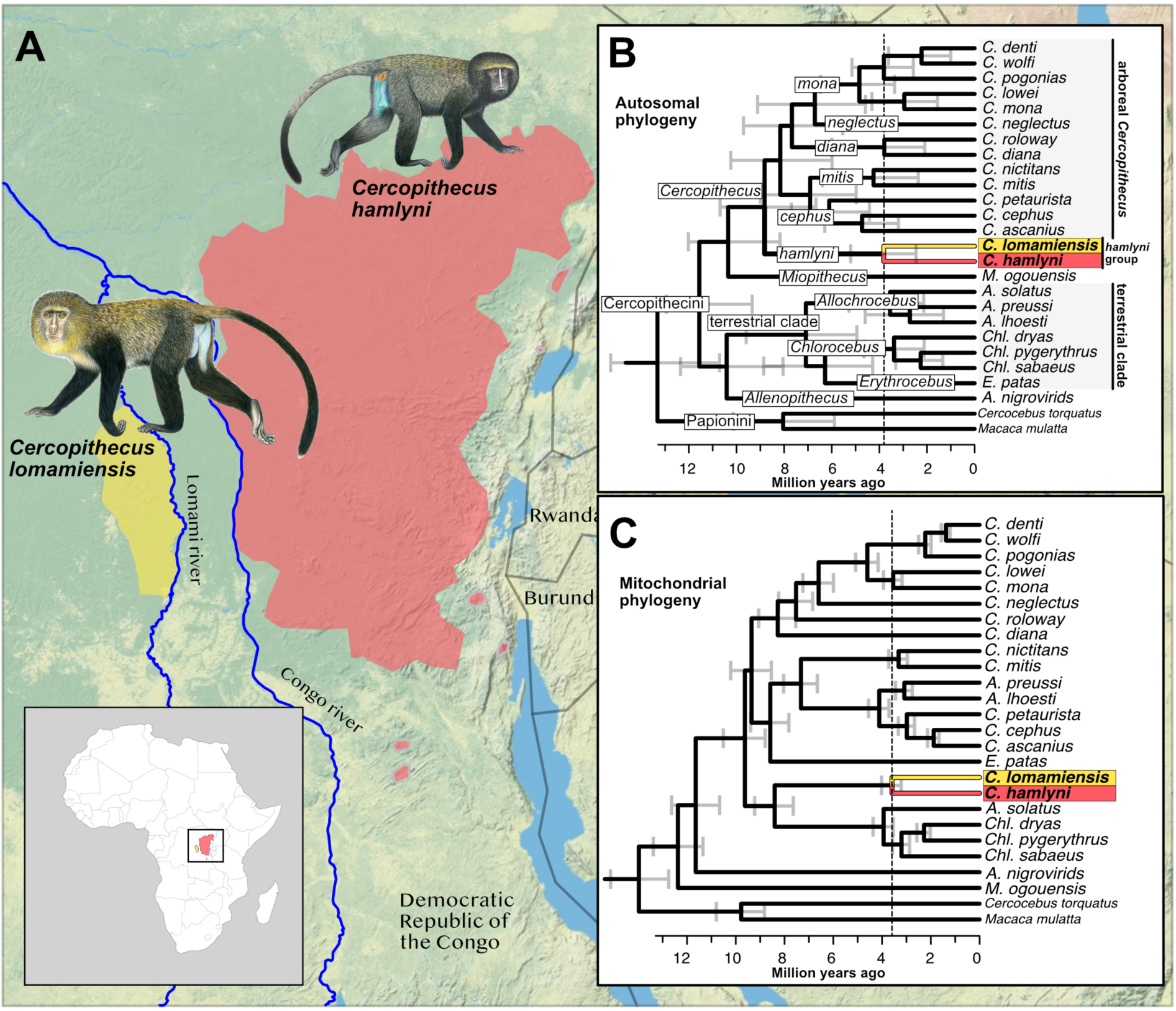
**A)** Distribution ranges of *C. lomamiensis* (yellow) and *C. hamlyni* (red), based on data from IUCN (Hart & Maisels 2020, Detwiler & Hart, 2020). **B)** Autosomal phylogeny with divergence date estimates for included species. Branch annotations show tribes (capitalized, non-italics), genera (capitalized, italics), and species groups (non-capitalized, italics). **C)** Mitochondrial phylogeny and divergence date estimates for the same species as in (B). Vertical dashed lines in (B) and (C) highlight the estimated split time between *C. lomamiensis* and *C. hamlyni*. Topologies were pruned to include a single sample per species prior to divergence date estimates, treating *C. albogularis* and *C. mitis* as conspecifics due to paraphyly (Jensen et al. 2024). Complete trees with branch support annotations are displayed in Figure S1. Primate illustrations copyright 2013 Stephen D. Nash/IUCN SSC Primate Specialist Group. Used with permission.

As a member of the speciose and predominantly arboreal *Cercopithecus* genus, the *hamlyni* group stands out with its predominantly terrestrial lifestyle (Arenson et al., 2020; Fournier, McPhee, et al., 2023). In this context, the *hamlyni* group resembles the more distantly related “terrestrial clade” guenons (genera *Allochrocebus*, *Chlorocebus* and *Erythrocebus*; Figure 1B; Jensen et al., 2023; Tosi et al., 2004). The number of transitions between terrestriality and arboreality among guenons have been long debated, as different phylogenetic markers reached different conclusions regarding the monophyly of this trait (Gebo & Sargis, 1994; Guschanski et al., 2013; Tosi et al., 2004). The most recent and comprehensive guenon phylogeny, based on whole genome sequencing data from 22 guenon species, however, confirm that the *hamlyni* group is nested within a clade consisting of the predominantly arboreal genera *Miopithecus* and *Cercopithecus* (Jensen et al., 2023). Thus, the transition to terrestriality in the *hamlyni* group likely occurred independently from that in the terrestrial clade, or constitute a retained ancestral trait (Arenson et al., 2020; Lo Bianco et al., 2017).

The guenons underwent an extensive diversification over the past ∼12 million years, and currently comprise more than 30 recognized species (Grubb et al., 2003; Jensen et al., 2023; Lo Bianco et al., 2017). Contemporary hybridization, also across divergent lineages, is not uncommon (Detwiler, 2019; Jong & Butynski, 2010), and Jensen et al. (2023) demonstrated that the guenon radiation was shaped by extensive hybridization and gene flow among ancestral lineages. Interspecific gene flow is a potent evolutionary force, with the ability to facilitate adaptation and diversification (Abbott et al., 2013), and can potentially explain traits that are incongruent with a bifurcating species tree, such as terrestrial locomotion in the *hamlyni* group and the terrestrial clade.

Here, we sequenced the genome of the holotype specimen *C. lomamiensis*, and by integrating it with recently sequenced *C. hamlyni* individuals (Jensen et al., 2024, 2023; Kuderna et al., 2023), we performed a comprehensive genomic analysis of this poorly known species group. We evaluated the genomic diversity of the recently described *C. lomamiensis* and its sister species *C. hamlyni* relative to other *Cercopithecus* species, and identified genes under positive selection in the *hamlyni* group ancestor. We also explored the role of gene flow in the evolution of *C. lomamiensis* and *C. hamlyni*, specifically investigating if introgression from the terrestrial clade may have facilitated a shift to terrestriality in the *hamlyni* group.

## Methods

### DNA library preparation and sequencing

DNA was extracted from the holotype specimen of *C. lomamiensis* (YPM MAM 14080; Hart et al., 2012) in the Primatology Lab at Florida Atlantic University (Institutional Biosafety Committee #2012-144, #2016-246) using the DNeasy Blood & Tissue Kit (Qiagen 69504; Germantown, MD). The sequencing library was prepared using the TruSeq PCRfree DNA preparation kit (Illumina Inc.), and sequenced on the NovaSeq 6000 platform (2 x 150 bp).

### Mapping and variant calling

Fifty-five publicly available genomes from 24 additional guenon species (Ayoola et al., 2020; Jensen et al., 2024, 2023; Kuderna et al., 2023; van der Valk et al., 2020) were compiled and processed together with the newly generated *C. lomamiensis* genome. These genomes contained two samples from *C. hamlyni*, the sister species of *C. lomamiensis*. We also included one genome each for *Macaca mulatta* (SAMN03264642) and *Cercocebus torquatus* as outgroups (Kuderna et al., 2023), which are members of Papionini, the sister tribe of the guenons. Raw sequencing reads were processed, mapped and genotyped following the Genome Analysis Toolkit (GATK) best practices pipeline (Caetano-Anolles, 2023). Remaining adapter content in the reads was marked with Picard/2.23.4 MarkIlluminaAdapters, and read group information added with Picard AddOrReplaceReadGroups. Next, the reads were mapped to the rhesus macaque (*Macaca mulatta*) reference genome (Mmul10, GCF_003339765.1) using the Burrows Wheeler aligner/0.7.17 (bwa mem, Li & Durbin, 2009), and sorted and deduplicated using Picard SortSam and Picard MarkDuplicates, respectively. Next, GATK/4.2 was used to call genotypes per sample with HaplotypeCaller, which were then combined across all samples and jointly genotyped with CombineGVCFs and GenotypeGVCFs, respectively. Insertions and deletions were excluded, and genotypes with read coverage below half or above twice the autosomal and X-chromosomal average at the sample level were masked on the autosomes and X chromosome, respectively. The Y chromosome was not included in downstream analyses. Sites with single nucleotide polymorphisms (SNPs) were additionally filtered with gatk VariantFiltration based on the recommended exclusion criteria (QD < 2.0, QUAL < 30.0, SOR > 3.0, FS > 60.0, MQ < 40.0, MQRankSum < -12.5, ReadPosRankSum < -8.0), and heterozygous genotypes with minor allele support < 0.25 were masked. Last, we used the SNPable regions pipeline (https://lh3lh3.users.sourceforge.net/snpable.shtml) to mask repetitive regions in the reference genome, using a Kmer size of 100 bp.

### Phylogeny and divergence times

We used IQ-TREE (v. 2.2.2.6-omp-mpi; (Minh et al., 2020) to construct maximum likelihood trees in genomic windows of 25 kb, separated by 500 kb across the reference autosomes. Windows with less than 10,000 genotyped sites were removed, and the remaining trees were used to estimate a species tree using ASTRAL (Zhang et al., 2018). Divergence date estimates were obtained using MCMCTree as implemented in paml/4.9j (Yang, 2007). We applied three fossil node calibrations, based on de Vries and Beck (2023): crown Cercopithecini (6.5-12.51 million years ago, Mya), the split between Cercopithecini and Papionini (6.5-15 Mya), and the split between *M. mulatta* and *Cercocebus torquatus* (5.33-12.51 Mya). Following the procedure described in Jensen et al. (2024), we randomly sampled 10 loci of 5 kb each from intergenic regions (≥ 10 kb from nearest gene), which were treated as different partitions in the analyses. Two independent MCMCTree analyses were conducted using the correlated clock model, and we discarded the first 10,000 iterations as burn-in and thereafter sampled every 100th iteration until 20,000 samples were collected. This procedure was then repeated across 10 independent sets of loci. After confirming convergence between the two runs within each locus set, all MCMCTree runs were merged and summarized jointly.

We assembled the mitochondrial genome of *C. lomamiensis* using MitoFinder (Allio et al., 2020), and combined it with previously assembled guenon mitogenomes (Jensen et al., 2024, 2023). The mitochondrial assemblies were annotated, processed and aligned as described in Jensen et al. (2023), and partitioned into 42 partitions: 1st, 2nd and 3rd codon position of each protein coding gene, the two ribosomal RNA, and all tRNA concatenated. The partitions were aligned using MAFFT (Katoh et al., 2019), and IQTree was used to perform a modeltest for each partition and estimate a maximum likelihood tree with 1,000 rapid bootstrap replicates. Mitochondrial divergence times were estimated with MCMCTree in two independent runs using the same settings as for the autosomal data, except that we reduced burn-in to 2,000 iterations and sampled every 50th iteration, for computational reasons.

### Demographic history and genetic diversity

We used beta-PSMC (Liu et al., 2022) to infer historical effective population sizes of *C. lomamiensis* and *C. hamlyni*. Beta-PSMC is based on the pairwise sequential Markovian coalescent method (PSMC; (Schiffels & Durbin, 2014), but suggested to have higher resolution particularly in recent times (Liu et al., 2022). To generate the input file used by beta-PSMC, we first converted the filtered genotype calls for each *hamlyni* group sample to fasta files with bcftools/1.20 consensus, which were subsequently converted into the psmcfa format using the fq2psmcfa script from the psmc utils suite (https://github.com/lh3/psmc). Beta-PSMC was run with recommended parameter settings (-p 20*1 -N25 -t15 -r5 -B5), with 20 bootstrap replicates. Since we observed a peak followed by a steep decline in the *C. lomamiensis* sample similar to artifacts that can be attributed to PSMC parameter settings (Hilgers et al., 2024, see results), we also ran the original PSMC software, modifying the parameter settings in recent time segments as recommended by Hilgers et al. (2024), with similar results.

Next, we used a custom Python script to estimate genome-wide heterozygosity, calculated as the number of heterozygous autosomal genotypes divided by the total number of called autosomal genotypes per sample. We also used Plink (Purcell et al., 2007) to identify autosomal runs of homozygosity (ROH) of at least 100 kb. To estimate the fraction of the genome contained in ROHs, we first estimated the total length of the genome accessible to the ROH analysis, following Meyermans et al. (2020). This was done by converting all heterozygous genotypes to homozygous calls, and running the same ROH identification on this artificial, completely homozygous genome. The fraction of each individual genome contained in ROHs was then calculated as the total length of identified ROHs in the real data, divided by the total length of ROHs identified in the artificial, homozygous genome.

To investigate the presence of mutation load in the *hamlyni* group lineage, we used the Variant Effect Predictor (McLaren et al., 2016) to annotate the SNP calls with predicted functional impacts, based on the Mmul_10 genome annotation. We considered only impacts inferred from canonical transcripts, and quantified mutation load for alleles predicted to have moderate or high impacts separately, by dividing the total number of derived alleles in each category with the total number of predicted synonymous derived alleles.

### Identifying genes under selection

We used the HyPhy/2.5.60 suite to explore signals of selection in the *hamlyni* group (Kosakovsky Pond et al., 2019). Transcripts for all protein coding genes in the Mmul_10 genome were extracted to in-frame sequence alignments using a custom Python script, choosing the longest transcript for genes with multiple transcripts. Signals of positive selection in *hamlyni* group lineages were inferred using HyPhy absrel (M. D. Smith et al., 2015). This tool models a scenario with a proportion of sites evolving under positive selection in a predefined set of “foreground branches”, and compares it to a null model with all sites evolving neutrally or under purifying selection. If the model with positive selection provides a significantly better fit to the data than the null model based on a likelihood ratio test (LRT, p < 0.05), this is indicative of positive selection in the foreground branches. For these analyses, we included a single sample per species from all *Cercopithecus* species and *M. ogouensis* (the arboreal clade), and tested for signatures of positive selection separately in the ancestral *hamlyni* group branch and the terminal branches of *C. lomamiensis* and *C. hamlyni*, based on the species tree topology. We removed genes containing premature stop codons, and ran two independent analyses of each gene/foreground branch combination, and considered a result as significant only if both runs suggested the model of selection as a significantly better fit than the null model.

### Gene flow

To test for signals of introgression we used Dsuite (Malinsky et al., 2021) to calculate the D-statistics (Durand et al., 2011; Green et al., 2010) between the *hamlyni* group and other guenon lineages. The D-statistics quantifies excess allele sharing between either of two sister lineages (P1 and P2) and a third lineage (P3), using an outgroup to polarize the ancestral allele. To explore if any introgression occurred into either *C. lomamiensis* or *C. hamlyni* after they diverged from each other, we estimated D-statistics with *C. lomamiensis* as P1, iterating through the two *C. hamlyni* samples as P2 and all other guenon species as P3. *Macaca mulatta* was always used as the outgroup. We also calculated the D-statistics with *C. lomamiensis* or *C. hamlyni* as P3, *mitis* group spp. as P2 and *cephus*, *mona*, *diana* or *neglectus* group spp. as P1 (event C in Jensen et al. 2023), to confirm that the gene flow from the *mitis* group occurred in the *hamlyni* group ancestor.

Next, we revisited the previously reported gene flow event from the terrestrial clade ancestor to the *Cercopithecus* ancestor (event B in Jensen et al., 2023), to investigate whether this could have facilitated adaptations to the terrestrial lifestyle of *C. hamlyni* and *C. lomamiensis*. Genome-wide D-statistics were calculated for all combinations of terrestrial clade spp. as P3, *Cercopithecus* spp. as P2 and *Miopithecus ogouensis* as P1. We also inferred the genomic landscape of introgression from the terrestrial clade into the *Cercopithecus* species, by estimating *f_dM_* in non-overlapping, sliding windows of 10 kb, along the reference genome, using Simon Martin’s ABBABABAwindows.py script (https://github.com/simonhmartin/genomics_general). The *f_dM_* statistic is related to the D-statistic and also quantifies excess allele sharing indicative of gene flow, but is more robust to the sparsity of data associated with small genomic regions (Martin et al., 2015). After excluding windows with less than 100 SNPs, we compared *f_dM_* along the genome estimated with terrestrial clade spp. as P3 (donor lineage), *M. ogouensis* as P1 (control lineage), and either the *hamlyni* group or arboreal *Cercopithecus* as P2 (recipient lineage). The motivation for this analysis was that, if gene flow occurred only into the *Cercopithecus* ancestor, *f_dM_* estimates obtained with either the *hamlyni* group or the *Cercopithecus* species as P2 would be highly correlated and show almost a 1:1 relationship. We also estimated *f_dM_* using the *hamlyni* group spp. as P2 and all other *Cercopithecus* samples as P1, to specifically investigate the presence and function of privately retained introgression from the terrestrial clade ancestor in the *hamlyni* group.

After an initial gene flow event, introgressed haplotypes that segregate in the recipient population are expected to rapidly decrease in length as recombination breaks them up. Since the introgression from the terrestrial clade into the *hamlyni* group is very old (∼10 million years; Jensen et al., 2023), we expect most introgressed haplotypes to be very short, possibly making them difficult to detect using the *f_dM_* statistic. To investigate the presence of short but potentially functionally important introgression/private sorting of terrestrial clade alleles in the *hamlyni* group, we identified convergent amino acid (AA) substitutions in the *hamlyni* group and the terrestrial clade. Protein coding gene transcripts were extracted from the genotype calls as described in the section “Identifying genes under selection”, and converted into AA sequences with a custom Python script. Sequences with more than 50% missing data or internal stop codons were excluded, as were genes where more than half of the samples in any of the two groups were filtered out (due to missingness or internal stop codons). We then identified sites that showed a fixed private AA allele in the *hamlyni* group and the terrestrial clade genera *Erythrocebus* and *Chlorocebus* relative to the arboreal *Cercopithecus* spp. and *M. ogouensis*. As above, we excluded *Allochrocebus* from this test as this genus likely received additional gene flow from *Cercopithecus* (Jensen et al., 2023).

## Results

### Sequencing, mapping and variant calling

We sequenced ∼500 million read pairs from the *C. lomamiensis* genome, resulting in an average autosomal mapping depth of 49 x on the Mmul_10 reference genome (Table S1). We included 57 previously sequenced guenon and outgroup genomes (Ayoola et al., 2020; Jensen et al., 2024, 2023; Kuderna et al., 2023; van der Valk et al., 2020), summing up to a total of 58 genomes representing 25 guenon species and two Papionini outgroup species (*Macaca mulatta* and *Cercocebus torquatus*, Table S1). Median autosomal mapped read depth across the data set was 36.3 (19.0 - 63.2), and we retrieved a median of 2.37 Gb filtered, genotypes per sample across the autosomes and X chromosome (2.13 - 2.49).

### Phylogeny and divergence times

We used ASTRAL to construct a multispecies coalescent-based tree, from 5,037 autosomal gene trees. Our analyses confirm the autosomal sister relationship between *C. lomamiensis* and *C. hamlyni* (Hart et al., 2012), with full local posterior probability (LPP=1; Figure 1B, Figure S1). We also successfully assembled the mitochondrial genome of *C. lomamiensis*. Previous analyses have shown that *C. hamlyni* experienced mitochondrial introgression from the ancestor of the terrestrial clade (Jensen et al., 2023), whereas the mitochondrial phylogenetic placement of *C. lomamiensis* was unknown. The mitochondrial phylogeny shows a sister relationship between *C. hamlyni* and *C. lomamiensis* with full bootstrap support (Figure 1C, Figure S1), confirming that the mtDNA introgression from the terrestrial clade occurred in the *hamlyni* group ancestor. We estimated the autosomal divergence time between *C. lomamiensis* and *C. hamlyni* to ca. 3.8 million years ago (Mya; 95 % highest posterior density [HPD] = 2.5-5.2 Mya; Figure 1B) and the mitochondrial divergence to ca. 3.6 Mya (95%HPD = 3.2 - 4.0 May, Figure 1C).

### Demographic history and genetic diversity

We explored the demographic history of the two species in the *hamlyni* group by estimating the ancestral effective population size (*N_e_*) through time, using beta-PSMC. The *N_e_* trajectories of the two species diverge ca. 3-4 MYA, in agreement with the MCMCTree divergence time estimates (Figure 2A). *Cercopithecus lomamiensis* shows two peaks in *N_e_*, one at around 500,000 years ago reaching an *N_e_* of ca. 300,000, and a second peak at around 50,000 years ago with estimated *N_e_* of ca. 600,000, followed by a steep decline in the more recent time segments. Peaks similar to the highest and most recent one detected in *C. lomamiensis* can be caused by erroneous parameter settings in PSMC-based methods (Hilgers et al., 2024). Although beta-PSMC is expected to be relatively robust to such false peaks, we also repeated this analysis using PSMC, adjusting the parameter settings following Hilgers et al. (2024), and recovered a highly similar trajectory (Figure S2). Nevertheless, we caution that processes other than actual population size increase, e.g. population structure, could cause this peak and subsequent decline (Mazet et al., 2016). Population structure and admixture are also plausible mechanisms behind the steep *N_e_* increase in one of the analyzed *C. hamlyni* genomes, which separates from the other conspecific genome at around 300,000 years ago, reaching an unrealistic *N_e_* of ∼10,000,000 (Figure 2A). This sample comes from a captive individual of unknown provenance (Kuderna et al., 2023), and it is possible that it derives from an intraspecifically admixed population, which can cause similar peaks in recent time (Mazet et al., 2016).

**Figure 2.**
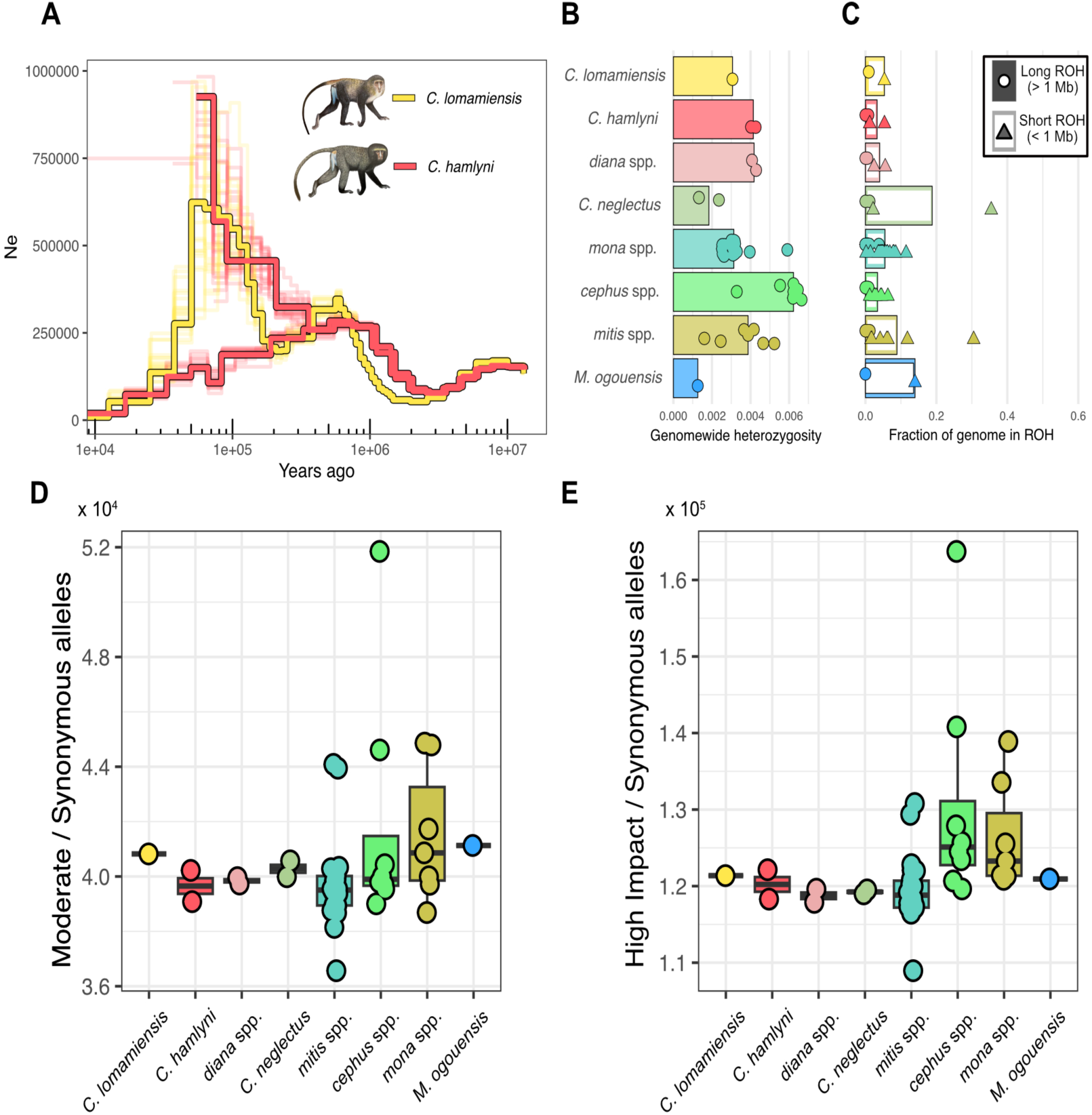
**A)** Effective population size (*N_e_*) through time for *C. lomamiensis* and the two *C. hamlyni* individuals, estimated with beta-PSMC. *N_e_* values above 1,000,000 were cut for visualization, full trajectory in Figure S2. **B, C)** Heterozygosity (B) and fraction of the genome in runs of homozygosity (ROH) (C) for *C. lomamiensis*, *C. hamlyni,* other included *Cercopithecus* species and *Miopithecus*, grouped by genus or species group. Each point represents one individual, and bars show the species group/genus average. In (C), circles and triangles show the proportion of the genome contained in long (>1Mb) and short (> 100Kb) runs of homozygosity, respectively, with averages per species group/genus represented as filled or open portion of the bars. **D, E)** Mutation load estimated as the number of derived alleles predicted to have moderate functional impact (D) or alleles predicted to have a high impact (E), divided by the number of synonymous derived alleles per individual. Primate illustrations copyright 2013 Stephen D. Nash/IUCN SSC Primate Specialist Group. Used with permission.

To evaluate the genomic diversity of *C. lomamiensis* and *C. hamlyni* in relation to other *Cercopithecus* taxa, we estimated the genome-wide heterozygosity, identified runs of homozygosity, and quantified genetic load in all *Cercopithecus* species. The *C. lomamiensis* individual showed lower genetic diversity than the two *C. hamlyni* individuals (Figure 2B), but the estimates of both species fall well within the range of other *Cercopithecus* species, and are comparable to, e.g., the more widespread and abundant members of the *mona* and *mitis* groups.

Runs of homozygosity (ROHs) are consecutive homozygous genomic segments that are identical by descent, and arise as a result of inbreeding (Ceballos et al., 2018). As recombination breaks down haplotypes over time, old inbreeding manifests as short ROHs, whereas longer ROHs indicate recent inbreeding. As expected given the relatively high heterozygosity, *C. lomamiensis* had a low ROH burden, comparable to most other *Cercopithecus* species (Figure 2C). Only 0.94 % of the genome was contained in long ROHs (> 1 Mb), and 4.4 % in short ROHs (100 kb - 1 Mb). A similarly low ROH burden was found in *C. hamlyni*. These results further support that neither *C. lomamiensis* nor *C. hamlyni* experienced recent inbreeding or severe historical bottlenecks.

In small populations, the efficacy of selection decreases whereas genetic drift increases. As a result, alleles that are deleterious and would otherwise be removed by selection may drift to high frequencies. To explore whether this could be a problem in *C. lomamiensis*, we estimated genetic load by counting alleles predicted to have a high, medium or low impact based on the Mmul_10 annotation (Figure 2D, 2E). Genetic load was highly similar across the *Cercopithecus* species including the *hamlyni* group, suggesting that the restricted distribution range of *C. lomamiensis* has not resulted in a detectable accumulation of deleterious alleles. We also estimated heterozygosity for each SNP category separately, and found that heterozygosity was higher in SNPs with moderate and high functional impact compared to synonymous mutations (Figure S3), suggesting that purifying selection is acting on high impact mutations, keeping them at low frequency.

### Adaptive evolution in the *hamlyni* species group

To explore genomic signatures of adaptive evolution in the *hamlyni* species group, we used HyPhy absrel to identify genes evolving under positive selection in the *hamlyni* ancestor, and the terminal branches of *C. lomamiensis* and *C. hamlyni*, respectively (Figure 3). After removing genes with internal stop codons, and genes with collapsed foreground branches (i.e. branch lengths of zero), we were able to test 12,569 genes for positive selection in the *hamlyni* ancestor, and 16,961 in the respective terminal branches. We found 111 genes for which the model with a proportion of sites evolving under positive selection in the *hamlyni* ancestor was a better fit to our data than a null model of neutral evolution (LRT, p < 0.05, Table S2). The number of genes with a significant signal of positive selection in *C. lomamiensis* and *C. hamlyni* were 45 and 33, respectively (Table S3, S4). Three genes, *TLR4*, *APOL6* and *LOC711523*, were identified as under positive selection in both *C. lomamiensis* and *C. hamlyni*.

**Figure 3.**
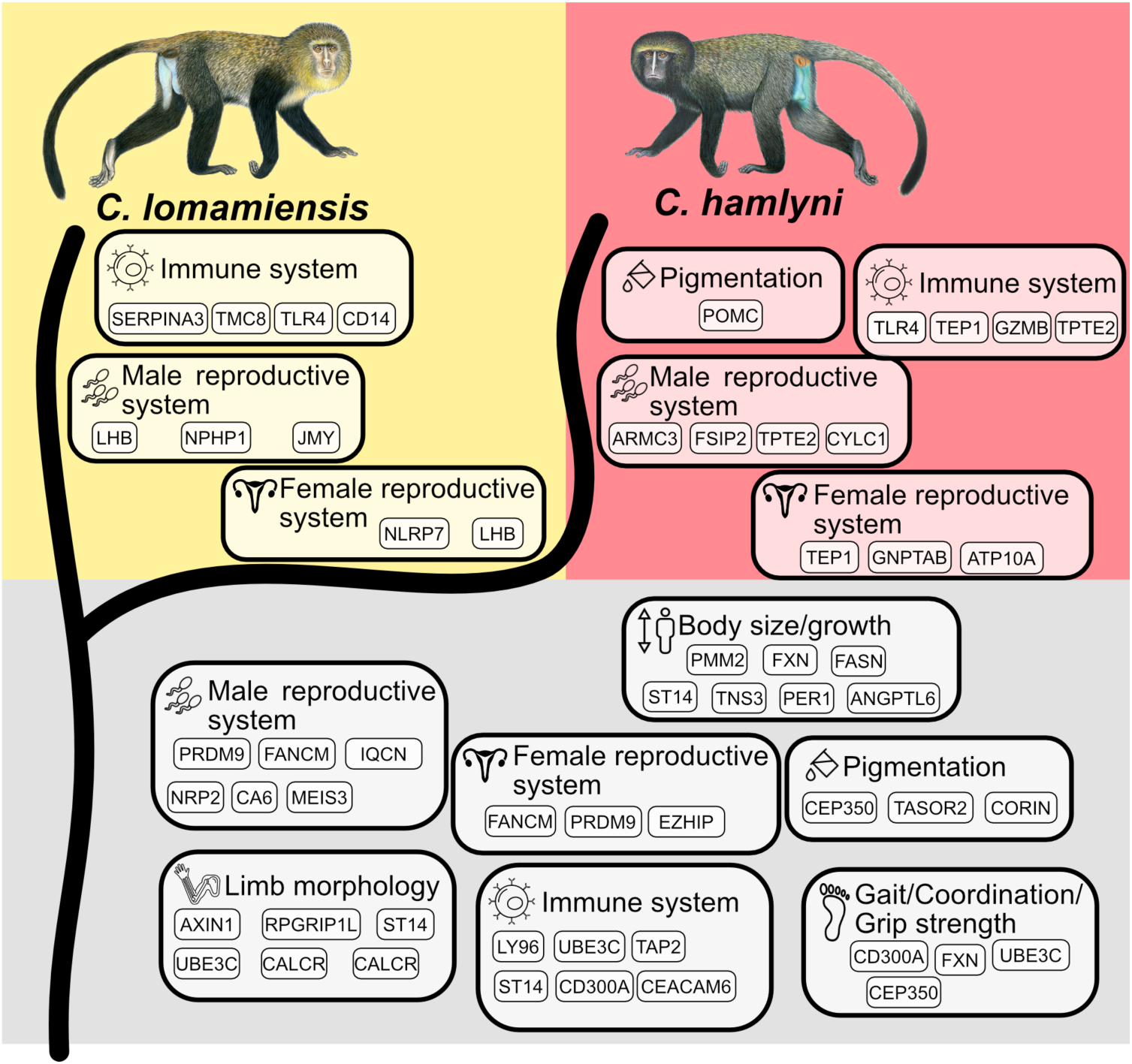
Examples of functions associated with genes identified as being under putative positive selection in the *hamlyni* group ancestor (along the basal lineage, gray background), *C. lomamiensis* (yellow background) or *C. hamlyni* (red background) terminal branches. Functional assignments were based on phenotype annotations from the mammalian phenotype database (C. L. Smith & Eppig, 2009). Note that these categories do not represent enrichments among genes under selection, and the genes under each category are examples. The complete set of phenotype annotations, and genes identified as being under positive selection, are presented in Table S6-S7 and Table S2-S4, respectively. Primate illustrations copyright 2013 Stephen D. Nash/IUCN SSC Primate Specialist Group. Used with permission.

We used the mammalian phenotype database to explore the functions of genes putatively under selection (C. L. Smith & Eppig, 2009). No mammalian phenotype (MP) or genotype ontology (GO) enrichments were found among genes under positive selection, neither in the *hamlyni* ancestor nor in the terminal *C. lomamiensis* or *C. hamlyni* branches (FDR > 0.05). In total, 854 MP terms were associated with the genes putatively under positive selection in the *hamlyni* ancestor (Table S5). These included, for example, functions related to the immune system, male and female reproductive traits, metabolism, and body size. We also highlight three genes annotated with phenotypes related to pigmentation (*CEP350*, *TASOR2*, and *CORIN*) as potentially important for pelage coloration evolution (Figure 3).

The genes with signatures of positive selection in the terminal branches of *C. lomamiensis* and *C. hamlyni* were associated with 380 and 884 phenotype terms, respectively (Table S6, Table S7). In both species, the most common terms were again related to, e.g., reproductive traits and immune functions (Figure 3). One gene involved in coat coloration, *POMC*, was found to be under positive selection in *C. hamlyni*. This gene has functions in melanogenesis (Millington, 2006), and could potentially be involved in the darker species-specific pelage color of *C. hamlyni* compared to *C. lomamiensis*.

### The role of ancestral gene flow in the *hamlyni* group evolution

Jensen et al. (2023) found that *Cercopithecus hamlyni* was involved in two gene flow events: One from the terrestrial clade ancestor into the *Cercopithecus* ancestor, and a second between the *mitis* group and *C. hamlyni*. Since the genome of *C. lomamiensis* was not previously available for analyses, we first estimated D-statistics to test if *C. lomamiensis* and *C. hamlyni* differed from each other in the patterns of gene flow with other guenon species. We found no or only negligible excess allele sharing between *C. lomamiensis* and other guenons, relative to *C. hamlyni* (Figure S4A). This confirms that the previously identified gene flow events occurred with the *hamlyni* group ancestor (Figure S4B), and suggest that no detectable, additional gene flow occurred that involved *C. hamlyni* or *C. lomamiensis* after the two species diverged.

Next, we focused specifically on the introgression event from the terrestrial clade ancestor, asking whether introgressed alleles may have facilitated adaptations to terrestriality in the *hamlyni* group ancestor. In line with Jensen et al. (2023), we found highly similar D-statistic estimates among all *Cercopithecus* species, including *C. hamlyni* and *C. lomamiensis*, when testing for excess allele sharing with the terrestrial clade relative to *M. ogouensis* (Figure 4A). Furthermore, a sliding window *f_dM_* analysis, with either the *hamlyni* group or other *Cercopithecus* species as recipient lineage (P2), revealed a very similar genomic landscape of introgression from the terrestrial clade (Figure 4B, Pearson’s r = 0.84, p < 0.001). Such similar genome-wide D-statistics and genomic landscapes of introgression strongly suggest that gene flow indeed predominantly occurred in the *Cercopithecus* ancestor.

**Figure 4.**
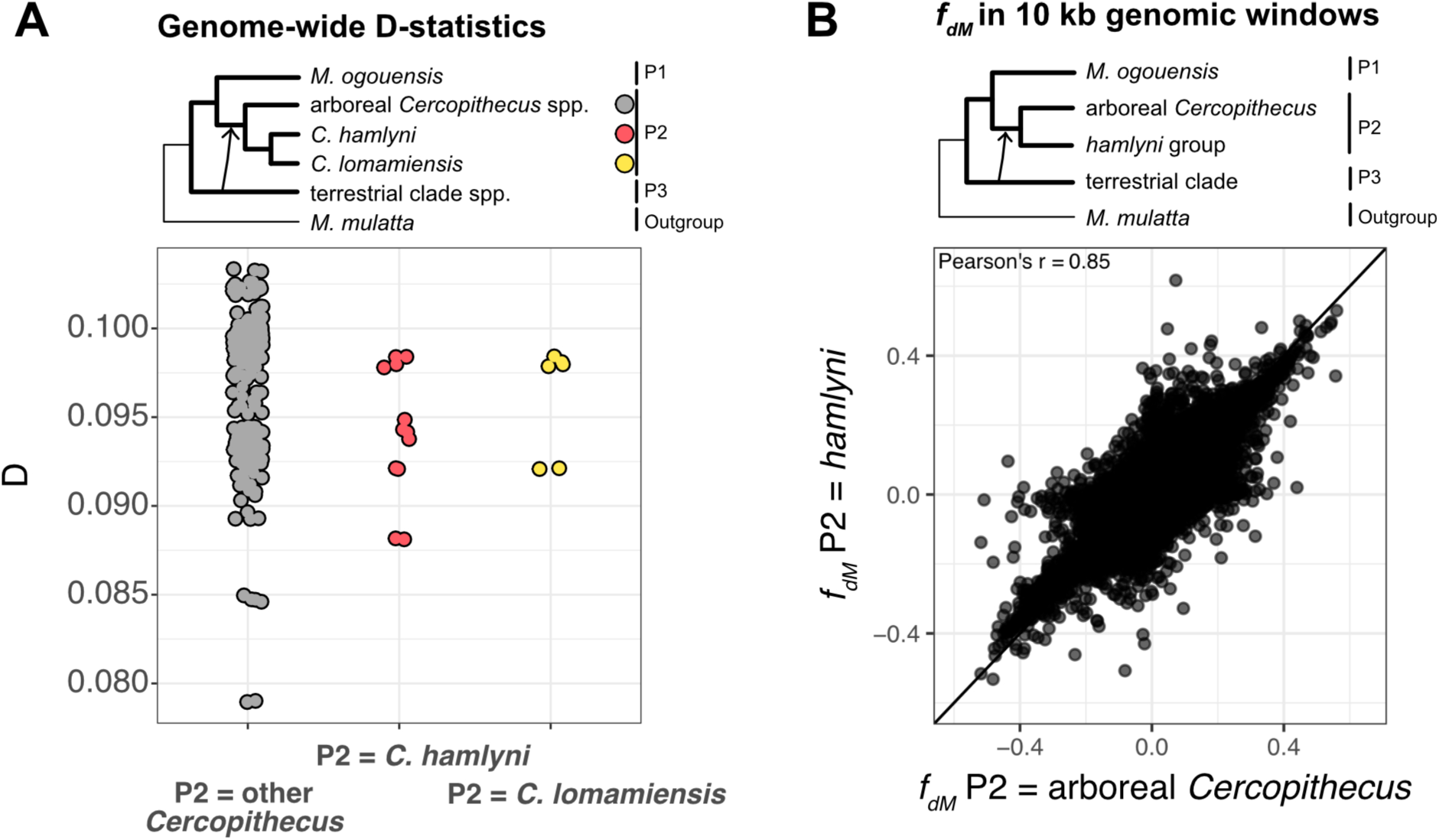
Excess allele sharing between the *Cercopithecus* spp. and the terrestrial clade. **A)** D-statistics results for tests with *M. ogouensis* as P1, *Cercopithecus* spp. as P2 and terrestrial clade spp. as P3. **B)** Excess allele sharing in genomic windows between terrestrial clade species and *hamlyni* group species (Y-axis) or other *Cercopithecus* species (X-axis). Each point represents one genomic window of 10 kb. The diagonal line shows 1-to-1 correspondence, i.e. the expected distribution of points under identical signals of introgression in the two lineages.

Nevertheless, the privately retained terrestrial clade-like mtDNA in the *hamlyni* group shows that either i) undetectably low levels of gene flow occurred from the terrestrial clade into the *hamlyni* group after it diverged from the other *Cercopithecus*, or ii) this mitochondrion - and thus potentially also other introgressed alleles - were differentially sorted among *Cercopithecus* lineages and retained only in the *hamlyni* group ancestor. To test for private retention of adaptive variants with functions potentially related to terrestriality in the *hamlyni* ancestor, we estimated excess allele sharing (*f_dM_*) between these lineages in 10 kb sliding windows. In line with lack of gene flow after the split between the *hamlyni* lineages and the other *Cercopithecus*, the *f_dM_* estimates were approximately normally distributed around a mean close to zero (mean *f_dM_* = 0.00047, Figure S5A). Furthermore, there was a strong heterogeneity in *f_dM_* estimates along the genome, without any clear peaks indicative of long introgressed genomic regions (Figure S5B).

Although our gene flow analyses suggest that private retention of introgression from the terrestrial clade in the *hamlyni* group was negligible at the genome-wide scale, functionally important haplotypes may still have been differentially sorted among *Cercopithecus* lineages. We therefore explored the gene content in genomic regions with strong signals of introgression from the terrestrial clade in the *hamlyni* lineage. We found 760 genes with coding sequences or upstream regions (< 10 kb) overlapping genomic windows with *f_dM_* estimates in the 99^th^ percentile. No MP or GO enrichments were found among these genes (FDR > 0.05), but several were annotated with phenotype terms that may be involved in adaptations to terrestrial locomotion (Table S8). These included both MP terms related to neurological or behavioral functions, e.g., ‘abnormal gait’ (30 genes), ‘decreased grip strength’ (30 genes), ‘impaired coordination’ (17 genes) and ‘abnormal locomotor behavior’ (16 genes), and terms related to morphological features, such as ‘short tibia’ (16 genes), ‘short femur’ (10 genes) and ‘short tail’ (4 genes). Although some of these genes may have been involved in terrestrial adaptations, only five of them showed a signature of positive selection in the *hamlyni* group ancestor (*CCDC7*, *LOC114673075*, *MUC16*, *SPHKAP* and *TNS3*), which is not a significant overlap (Fisher’s exact test, p = 0.4, Figure 5). We hence find no support for a prominent role of introgression from the terrestrial clade ancestor in the evolution of terrestriality in the *hamlyni* group.

**Figure 5.**
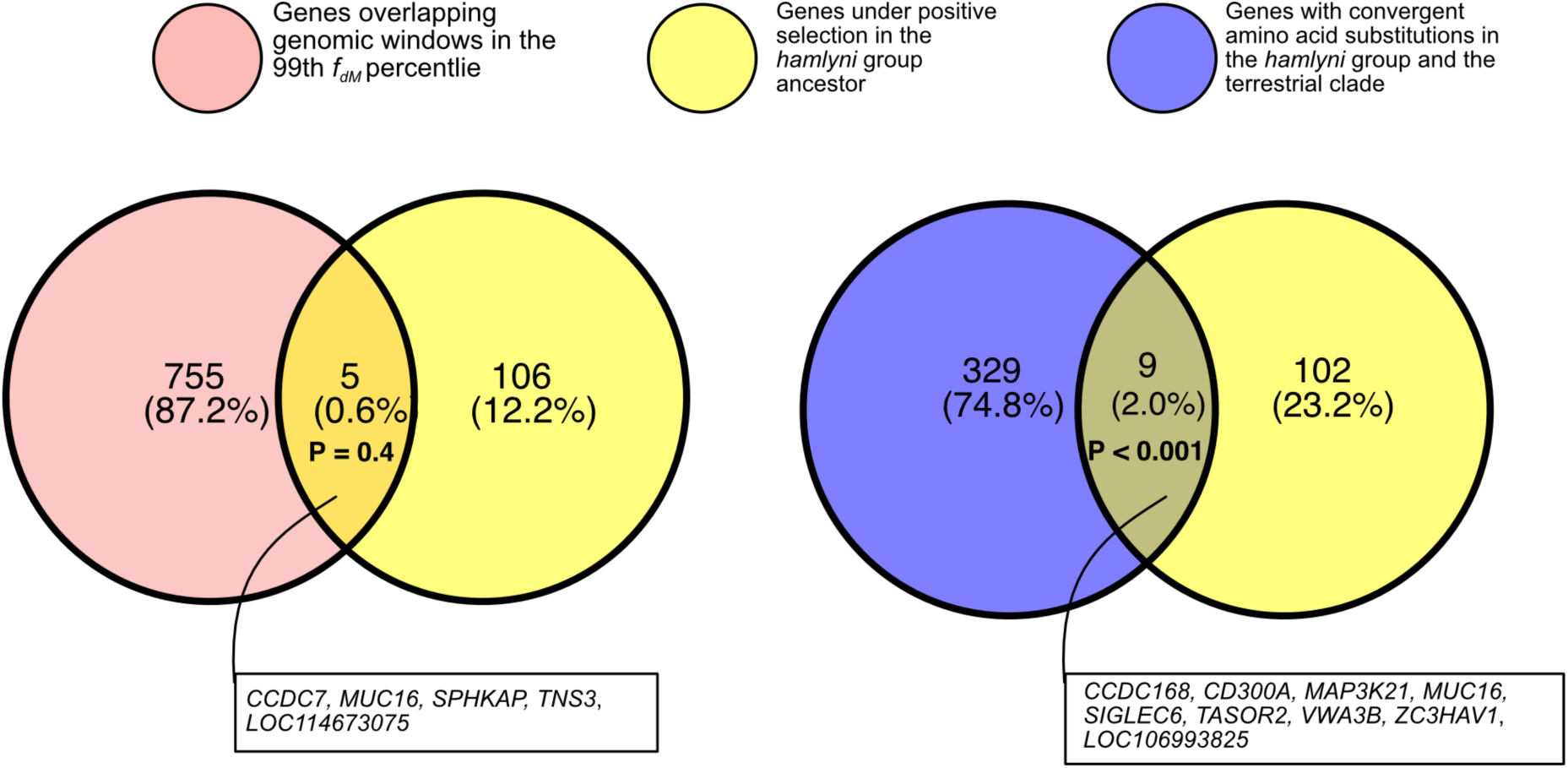
Overlap between genes with signals of positive selection in the *hamlyni* ancestral lineage and genes overlapping genomic regions with strong signals of introgression from the terrestrial clade (*f_dM_* ≥ 99^th^ percentile, left), and between positively selected genes and genes with convergent amino acid substitutions in the *hamlyni* group and the tererrestrial clade lineage.

Since the introgression from the terrestrial clade occurred prior to the radiation within the genus *Cercopithecus* ca. 10 million years ago, introgressed haplotypes may be very short and possibly difficult to detect using window-based analyses, such as the *f_dM_* statistic. To investigate the presence of introgression or differential retention of short but functionally important alleles, we identified convergent amino acid (AA) substitutions along the terrestrial clade and the *hamlyni* group ancestors, relative to the arboreal *Cercopithecus* spp. and *M. ogouensis*. The numbers of convergent AA substitutions were sparse overall, and similar in genes under positive selection in the *hamlyni* ancestor and other genes (Figure S6). A total of 338 genes contained at least 1 convergent AA substitution, with the maximum number of thirteen such substitutions found in the *CHIA* gene. This gene has been associated with insectivory in primates (Janiak et al., 2018), and could potentially indicate some dietary convergence in the *hamlyni* group and the terrestrial clade, but further research is needed to explore this. In contrast to the window-based *f_dM_* analysis above, we found a significant overlap between genes under positive selection in the *hamlyni* group and those with convergent AA substitutions with the terrestrial clade (n = 9, p < 0.001, Figure 5). This may suggest some degree of molecular convergence between the terrestrial clade and the *hamlyni* group in these genes, even if generally the results are more in line with a scenario of independent adaptations to terrestriality in the respective lineage.

## Discussion

In this study, we sequenced the genome of the elusive lesula, *Cercopithecus lomamiensis*, an African primate species first described to the scientific community just over a decade ago (Hart et al., 2012). Our results confirm the sister relationship between *C. lomamiensis* and *C. hamlyni*, as previously suggested based on morphology, vocalizations and sex chromosome markers (Arenson et al., 2020; Hart et al., 2012). We estimate that *C. lomamiensis* and *C. hamlyni* diverged ca. 3-4 million years ago (Mya), which is on par with (but slightly older than) previous estimates based on X-linked markers (∼2.8 Mya; Hart et al., 2012). The use of recently curated fossil calibrations (De Vries & Beck, 2023) in combination with more data is likely the cause of this discrepancy.

Despite the restricted distribution range of *C. lomamiensis*, we found no indications of inbreeding, reduced genetic diversity or increased mutation load. Taken at face value, this suggests that there is no imminent risk to the species survival from genetic factors. The population of *C. lomamiensis* is believed to be relatively dense, with estimates exceeding 10,000 individuals (Detwiler & Hart, 2020; Hart et al., 2012), which is likely sufficient to prevent inbreeding. The genetic diversity of *C. hamlyni* was marginally higher than *C. lomamiensis*, and both species fall well within the range of other *Cercopithecus* spp., which is one of the most genetically diverse primate genera (Kuderna et al., 2023). Nevertheless, both *C. lomamiensis* and *C. hamlyni* are heavily hunted (Fournier, Graefen, et al., 2023), increasingly threatened by habitat degradation in parts of their range, and their populations are thought to be declining (Detwiler & Hart, 2020; Hart & Maisels, 2020). Our demographic analyses further suggest population declines in both species in the past ∼50,000-100,000 years, although we caution that this could be caused by population structure (Mazet et al., 2016). Temporal genomic studies are needed to fully understand how genetic diversity, inbreeding and mutation load are changing through time as a result of population declines (Díez-Del-Molino et al., 2018). Previous analyses have suggested that genetically diverse species experiencing rapid population declines suffer stronger genomic consequences than populations that have a long history of inbreeding (van der Valk, de Manuel, et al., 2019; van der Valk, Díez-del-Molino, et al., 2019). Therefore, close monitoring of population trends and genetic status will be important to ensure that viable populations are maintained. This work provides a comparative context for future studies, which can aid inferences on contemporary demographic processes.

We found signals of positive selection along the ancestral *hamlyni* group branch in 111 genes, with functions related to several potentially important traits. For example, guenons show a remarkable, species-specific diversity in pelage coloration, thought to be important for mate choice (Allen et al., 2014). In this context, we highlight the *CORIN* gene, which was putatively under positive selection in the *hamlyni* ancestor. In mice, this gene was identified as a regulator of the agouti pathway (Enshell-Seijffers et al., 2008), a key player in coloration across mammals (Sørensen et al., 2023) and birds (Robic et al., 2019; Semenov et al., 2021). The *CORIN* gene has also been implicated in pelage coloration variation in tigers (Xu et al., 2017). We also found genes involved in immune functions, which is expected given that pathogens are a strong selective force in most organisms, and have been shown to mediate adaptive evolution of immune genes also in guenons (Svardal et al., 2017). Guenons are natural hosts of the simian immunodeficiency virus (SIV), a close relative of the human immunodeficiency virus (HIV), which likely exerted a strong selective pressure. Indeed, in addition to genes annotated with immune related MP terms (Figure 3, Table S2-S4), several genes under positive selection in the *hamlyni* ancestor are reported to be involved specifically in the immune response to SIV. For example, expressions of *OAS2* is upregulated in the acute phase after SIV infection (Hosseini et al., 2015), the antiviral *ZC3HAV1* is an important inhibitor during SIV/HIV infection (Kmiec et al., 2020), and variation in the *TAP2* gene has been associated with HIV susceptibility in humans (Detels et al., 1996).

The phenotypic differences between *C. lomamiensis* and *C. hamlyni* are subtle, but they show divergence in, e.g., vocalization, skeletal morphology and pelage coloration (Arenson et al., 2020; Hart et al., 2012). In *C. hamlyni*, we found signatures of positive selection in, e.g., the *POMC* gene, which is involved in skin and hair pigmentation in humans (Millington, 2006). Although population level sampling of both species would allow for more detailed inferences of diversifying adaptive evolution in the *hamlyni* group through the detection of selective sweeps, both in coding regions and regulatory elements, our results provide valuable first insights into their genomic adaptations.

Both *hamlyni* group species have several anatomical features, mostly related to limb morphology, that likely represent adaptations to terrestriality, a trait that differentiate them from other *Cercopithecus* spp. which are generally arboreal (Arenson et al., 2020; Gebo & Sargis, 1994). Among the genes under positive selection in the *hamlyni* group ancestor, several were also annotated with phenotypes that could be related to a terrestrial lifestyle. The *CALCR* gene, for example, was annotated with the term “short femur” on the mammalian phenotype database (C. L. Smith & Eppig, 2009), and femur morphology is indeed one of the features that distinguished the *hamlyni* group from the arboreal *Cercopithecus* (Arenson et al., 2020). Genes annotated with phenotype functions such as limb morphology (e.g., *RPGRIP1L*, *ST14*, *AXIN1*, *UBE3C*), gait (*UBE3C*, *FXN*, *CD300A*) and grip strength (*CEP350*, *FXN*, *CD300A*), also provide potential candidates for adaptations to a terrestrial habitat. However, this remains speculative at this point, as the phenotypic effects of these genes in the *hamlyni* group are unknown.

The paraphyly of terrestriality as a trait is indisputable based on updated guenon phylogenies, as the terrestrial *hamlyni* group is nested within the arboreal *Miopithecus* and *Cercopithecus* genera to the exclusion of the terrestrial guenon clade containing the genera *Chlorocebus*, *Allochrocebus,* and *Erythrocebus* (Arenson et al., 2020; Jensen et al., 2023). Since the *Cercopithecus* ancestor received gene flow from the terrestrial clade ancestor, we investigated a potential role of introgression in the transitions between arboreality and terrestriality among guenons. Our results suggest that, despite a privately retained mtDNA, no detectable nuclear introgression specific to the *hamlyni* lineage occurred from the terrestrial clade ancestor.

Although early studies suggested that the guenon ancestor was likely arboreal (Tosi et al., 2004), pointing to multiple transitions to terrestriality, recent analyses based on revised phylogenies suggest (semi)terrestriality as the more likely ancestral state (Arenson et al., 2020). Under this scenario, the terrestriality of the *hamlyni* lineage may be equally parsimoniously inferred as a derived or retained ancestral state (Arenson et al., 2020).

Transitions to terrestriality have been rare among primate lineages (Estrada & Marshall, 2024), and are generally associated with several anatomical features such as short tail, long limbs and short digits (Elton et al., 2016; Garber, 2007). It is therefore likely that coding or regulatory changes in genes that shape such anatomical features may be expected in terrestrial primates.

Furthermore, this presumably makes terrestriality a highly polygenic trait: For example, ∼650 genes have been annotated with phenotypes related to limb morphology, ∼150 with digit morphology, and ∼300 with tail morphology in the mammalian phenotype database (C. L. Smith & Eppig, 2009). Although *C. lomamiensis* and *C. hamlyni* have morphological features that likely represent adaptations to a life on the ground, the differences to other, arboreal *Cercopithecus* are subtle (Arenson et al., 2020). Hence, the genomic components of these putative adaptations may be difficult to identify.

Overall, this study offers new insights into the genetic status and evolutionary history of the *Cercopithecus hamlyni* group lineage. We provide a first glance into their genomic signatures of selection, including potential adaptations to a terrestrial lifestyle. Although the genetic basis of this complex trait is not well characterized, multiple transitions between arboreality and terrestriality in guenons make them a suitable system for more studies on this topic in the future. We find that both *C. lomamiensis* and *C. hamlyni* currently retain genetic diversity on par with other guenon species. Although they do not appear to be threatened by genetic factors, the declining population trends led to their classification as vulnerable by the IUCN. The genomic resources from the type specimen provide a crucial reference for future research, enabling better-informed conservation strategies, deeper insights into evolutionary history, and potential tools for combating illegal wildlife trade (van der Valk & Dalen, 2024). To enhance protection of lesula in the buffer zone, conservation plans include implementing alternative livelihood activities for local communities and conducting targeted surveys. Additionally, there is a consideration of documenting traditional practices, including cultural prohibitions, to integrate them into more effective natural resource management approaches. This integration of genomic data with field-based conservation efforts represents an important model for how holistic approaches can benefit threatened species.

## Acknowledgements

We thank Eric Sargis and Julia Arenson for helpful comments on an earlier version of this manuscript, and Stacy-Anne Parke for discussions. We thank Terese and John Hart and the TL2 Project for their work on lesula conservation and research, whose efforts have enabled additional studies including this genomic analysis. We also acknowledge Radar Nishuli for his support of scientific research on primates in the region. Sequencing was performed by the SNP&SEQ Technology Platform in Uppsala. The facility is part of the National Genomics Infrastructure (NGI) Sweden and Science for Life Laboratory. The SNP&SEQ Platform is also supported by the Swedish Research Council and the Knut and Alice Wallenberg Foundation. The computations and data handling were enabled by resources in projects NAISS 2023/6-340 and NAISS 2023/5-506, provided by the Swedish National Infrastructure for Computing (SNIC) at Uppsala University (UPPMAX), funded by the Swedish Research Council through grant agreement no. 2022-06725. The project was supported by the Swedish Research Council VR (2020-03398) grant to K.G., Zoologiska Stiftelse grants to A.J., Margot Marsh Biodiversity Foundation and FAU Foundation, Inc. to K.M.D.

## Author contributions

Conceptualization: A.J., K.G., and K.M.D.; Methodology and analyses: A.J.; Sample acquisition: K.M.D.; Lab work: E.R.H and K.M.D.; Validation of conservation implications: K.B.; Writing of manuscript: A.J. and K.G. with contributions from K.M.D. All authors reviewed and approved the manuscript.

## Data availability

Whole-genome sequencing data from *C. lomamiensis* is available on European Nucleotide Archive under accession number [TBA, (data will be released upon acceptance)]. Accession codes to publicly available data used in this study are listed in Table S1.

## Code availability

Custom scripts used for analyses this manuscript are available on https://github.com/axeljen/lesula_manuscript.

