## Supplementary figures for "Holotype genome of the lesula provides insights into demography and evolution of a threatened primate lineage"

Jensen et al.

Supplementary Figures S1-S6

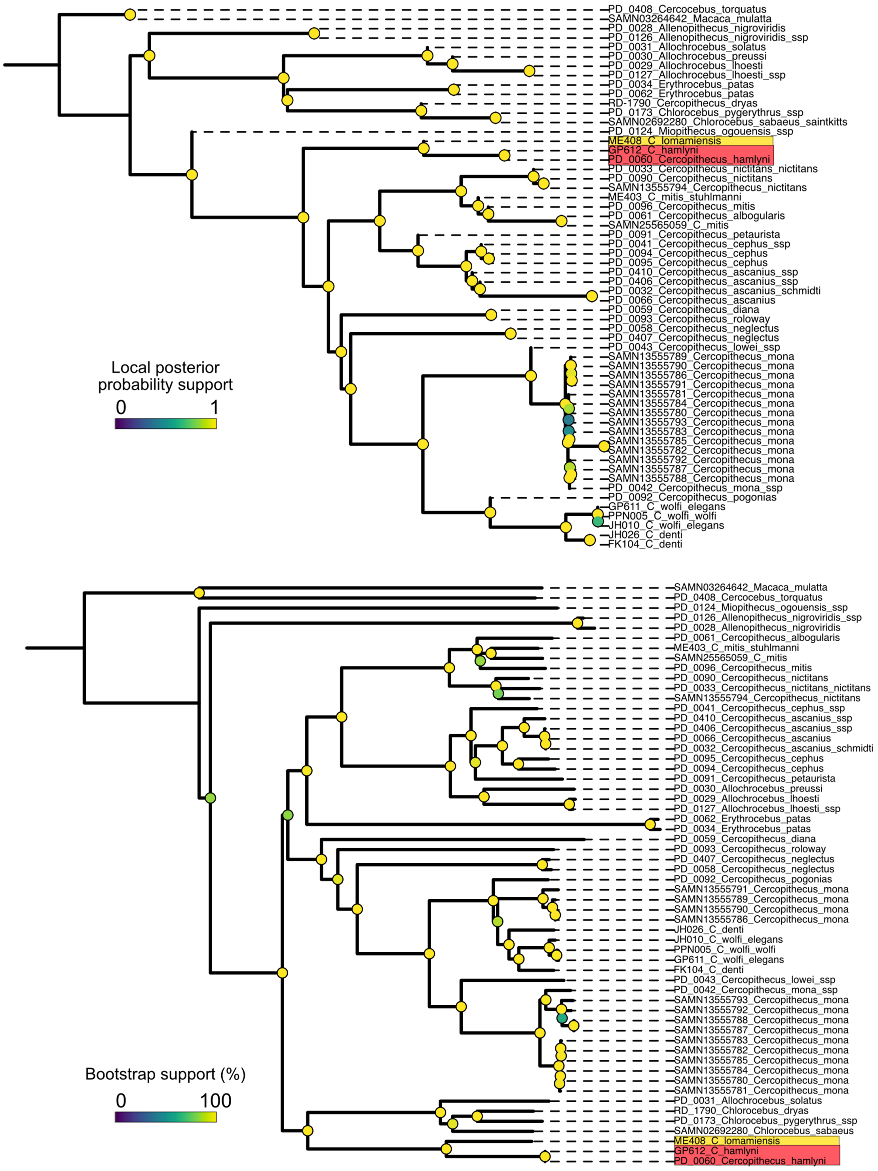


**Figure S1**. Autosomal coalescent-based tree estimated with ASTRAL (top) and mitochondrial maximum likelihood tree estimated with IQTree (bottom) with all samples included.

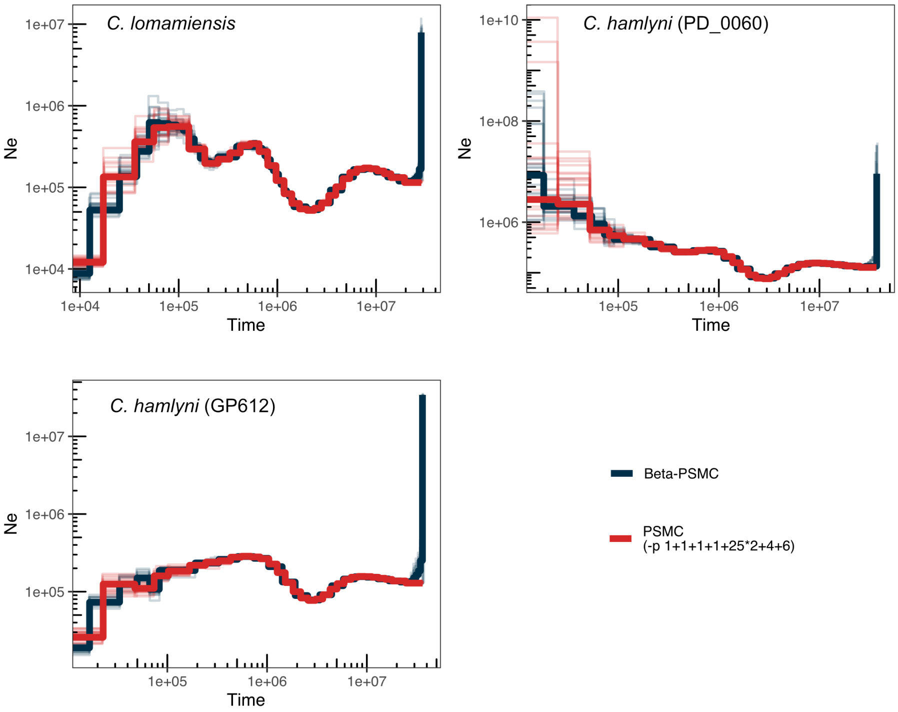


**Figure S2.** Effective population size through time estimated with beta-PSMC and PSMC, with the latter using parameter settings shown to decrease biases in recent times (Hilgers et al., 2024).

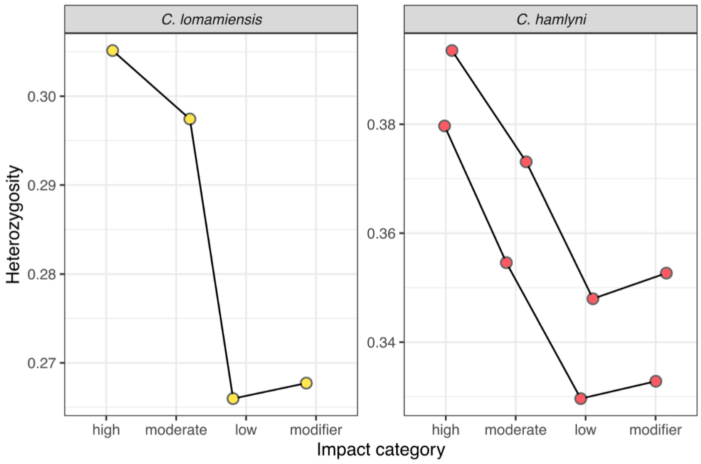


**Figure S3.** Heterozygosity of SNPs with different predicted impacts in *C. lomamiensis* and *C. hamlyni*.

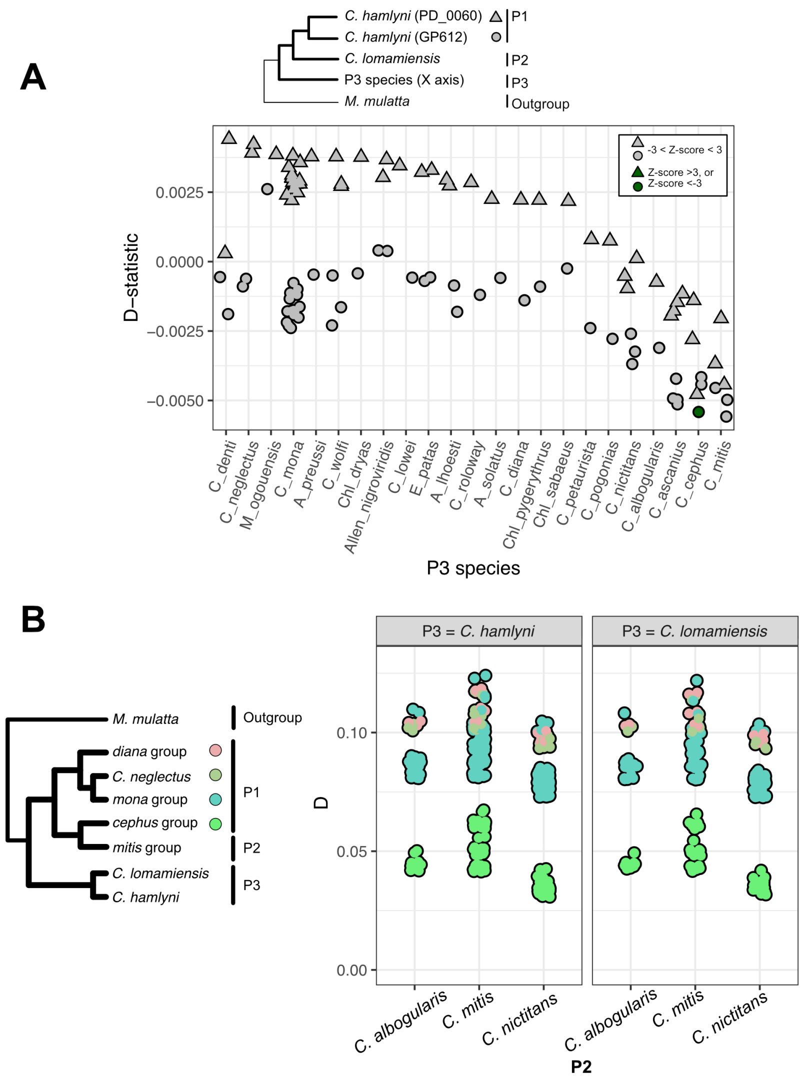


**Figure S4.** A) Excess allele sharing (D-statistics) between *C. lomamiensis* and all other guenon samples, relative to the two *C. hamlyni* samples (circles and triangles). Green circles/triangles indicate estimates that deviate significantly from 0 (Z-score > 3/Z-score < -3). B) Excess allele sharing (D-statistics) between *C. hamlyni/C. lomamiensis* and *mitis* group lineages. Highly similar estimates confirm that the gene flow between *C. mitis* and the *hamlyni* group occurred prior to the split between *C. hamlyni* and *C. lomamiensis*.

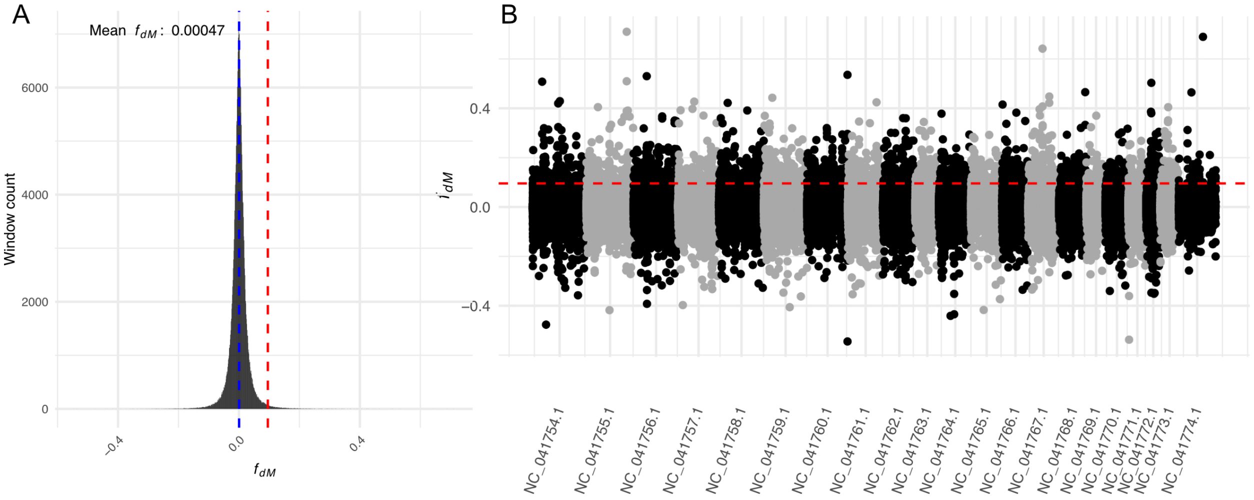


**Figure S5.** Genomic landscape of shared excess allele sharing between the *hamlyni* lineage and the terrestrial clade, relative to other *Cercopithecus* species (P1: *Cercopithecus* spp., P2: *hamlyni* group spp., P3: terrestrial clade spp., Outgroup: *M. mulatta*). A) Shows the distribution of $f_{dM}$ estimates in 10 kb windows, with the blue dashed line highlighting the genome-wide mean, and the red dashed line the 99$^{th}$ percentile. B) shows $f_{dM}$ estimates along the reference genome, with the dashed line highlighting the 99$^{th}$ percentile.

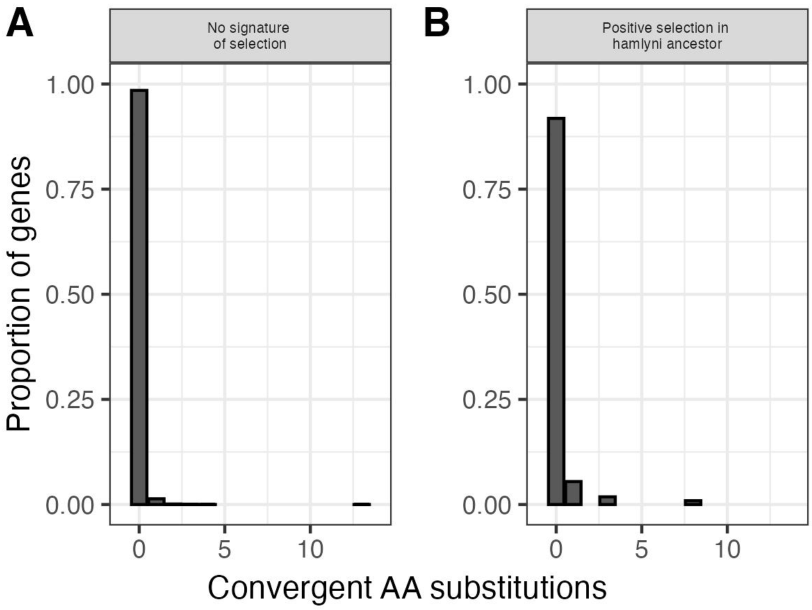


**Figure S6**. Proportion of genes with *X* number of convergent AA substitutions between the terrestrial clade linage and the *hamlyni* group, in (A) genes without any signature of selection and (B) genes with signatures of positive selection in the *hamlyni* ancestor, based on HyPhy analyses.
